## Supplementary Figures for "Divergent mechanisms of steroid inhibition in the human ρ1 GABA_A_ receptor"

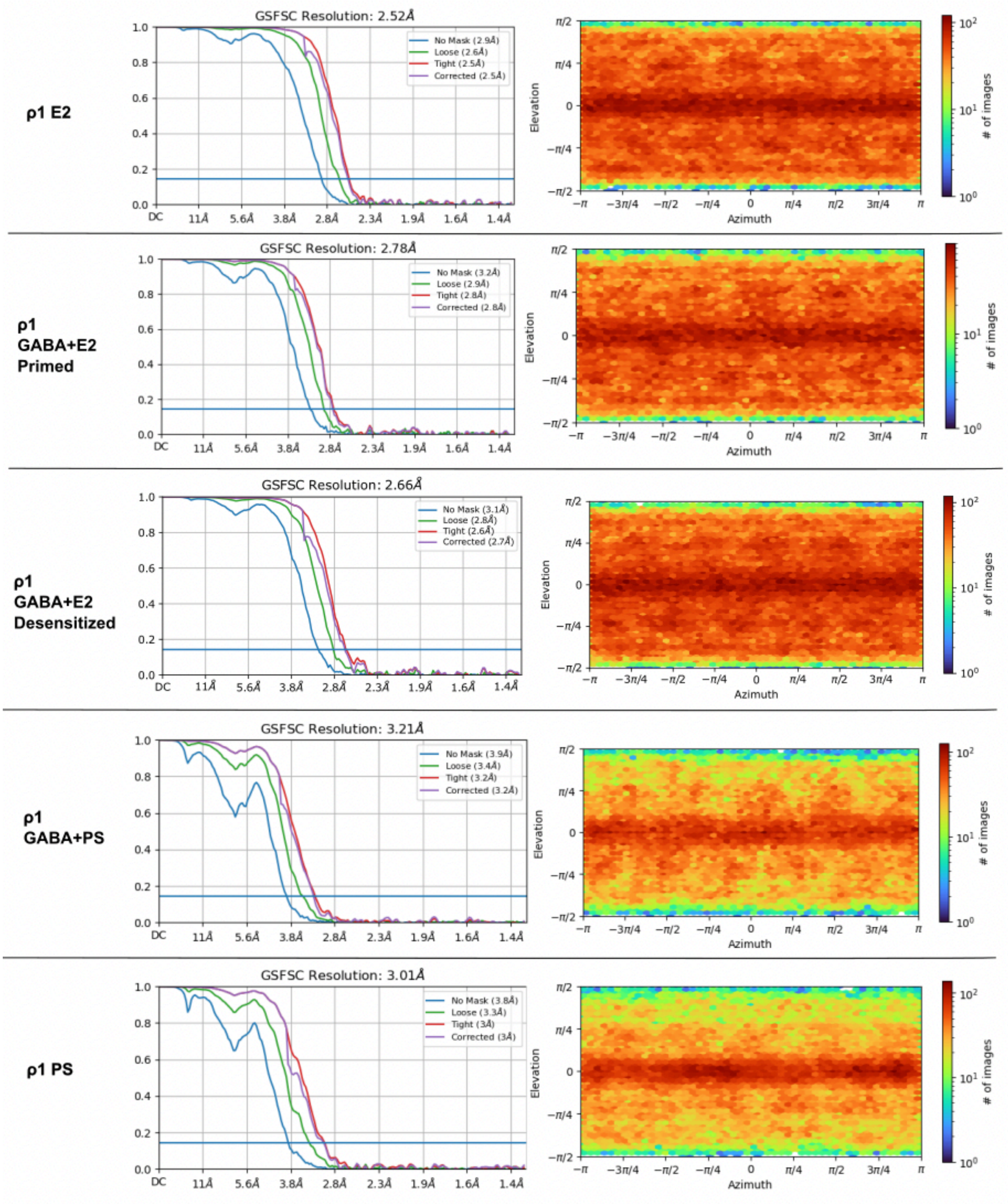


**Figure S1. FSC curves and angular distribution of the cryo-EM maps**

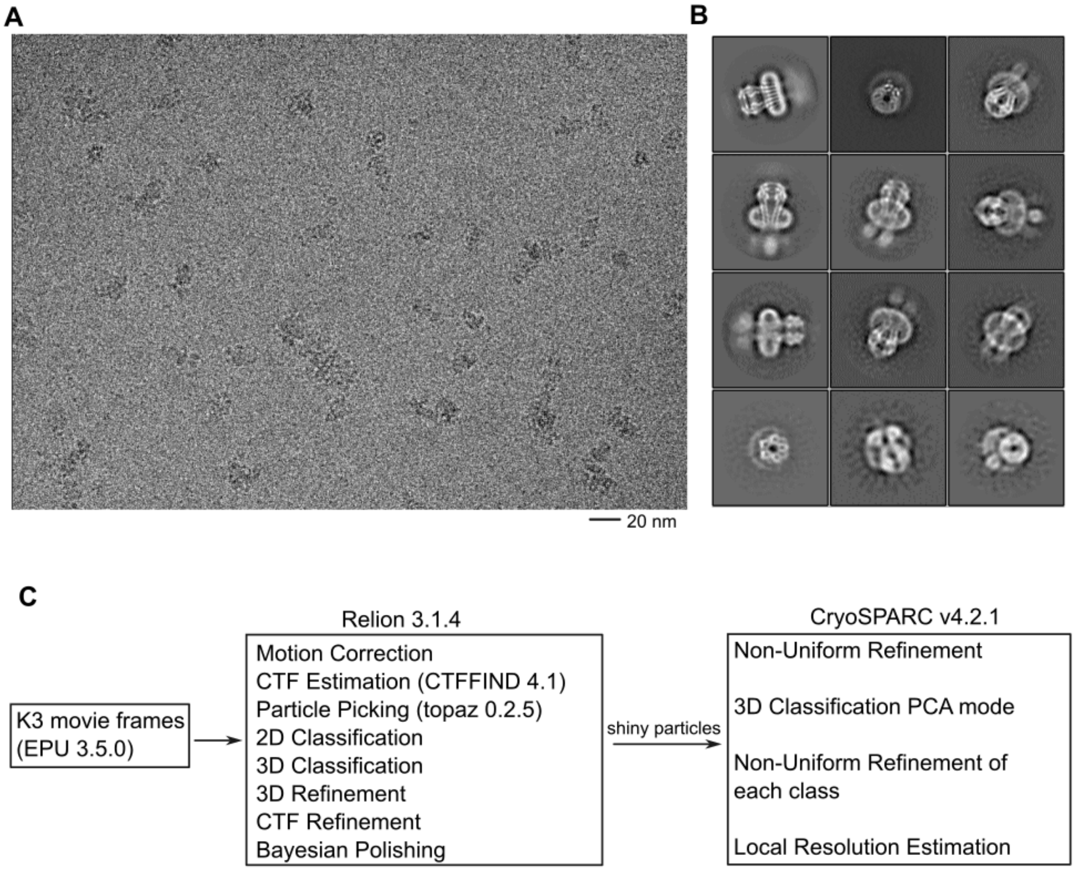


**Figure S2. Processing pipelines for ρ1-EM structures**

(A) Representative cryo-EM image from the ρ1-EM/E2 dataset.

(B) Representative 2D classification images of ρ1-EM/E2 dataset.

(C) Cryo-EM data processing workflow for all datasets reported in this work.

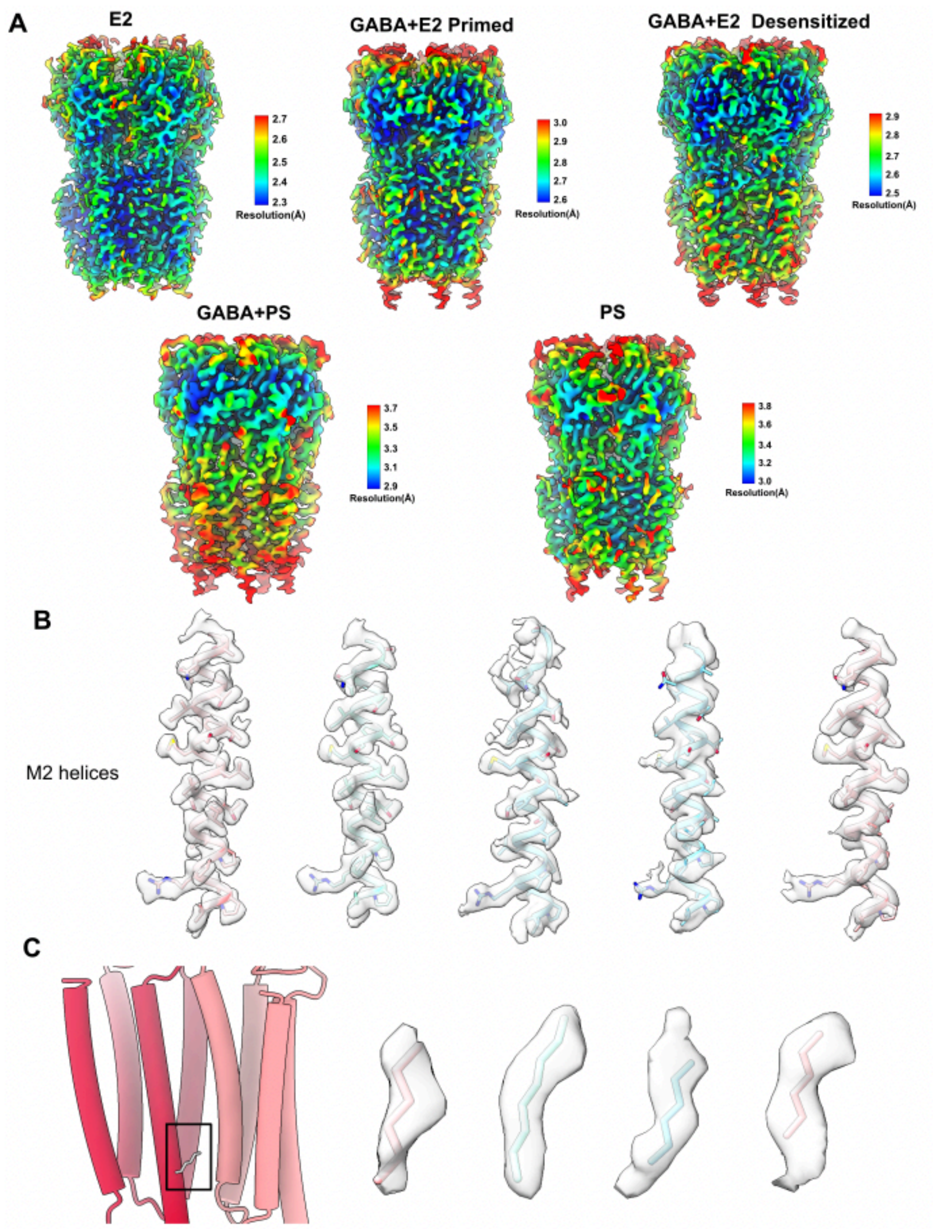


**Figure S3. Local resolution and representative densities in ρ1-EM structures**

(A) Cryo-EM maps colored by local resolution.

(B) Densities and models of M2 helices from all structures reported in this study, determined (left to right) in the presence of E2, GABA+E2 in the primed state, GABA+E2 in the desensitized state, GABA+PS, and PS alone.

(C) Apparent lipid at the M3-M1(-) interface of several ρ1-EM structures, corresponding to the allopregnanolone binding site in the α1β2γ2 $GABA_A$ receptor. Leftmost cartoon shows a TMD interface between two subunits in the structure with E2, with the relevant lipid as sticks. Remaining models (left to right) show corresponding densities in structures determined with E2, GABA+E2 in the primed state, GABA+E2 in the desensitized state, and PS alone, enlarged and rendered transparent, with atoms modeled as segments of phospholipid tails. No clear lipid density was observed in the ρ1 GABA+PS structure.

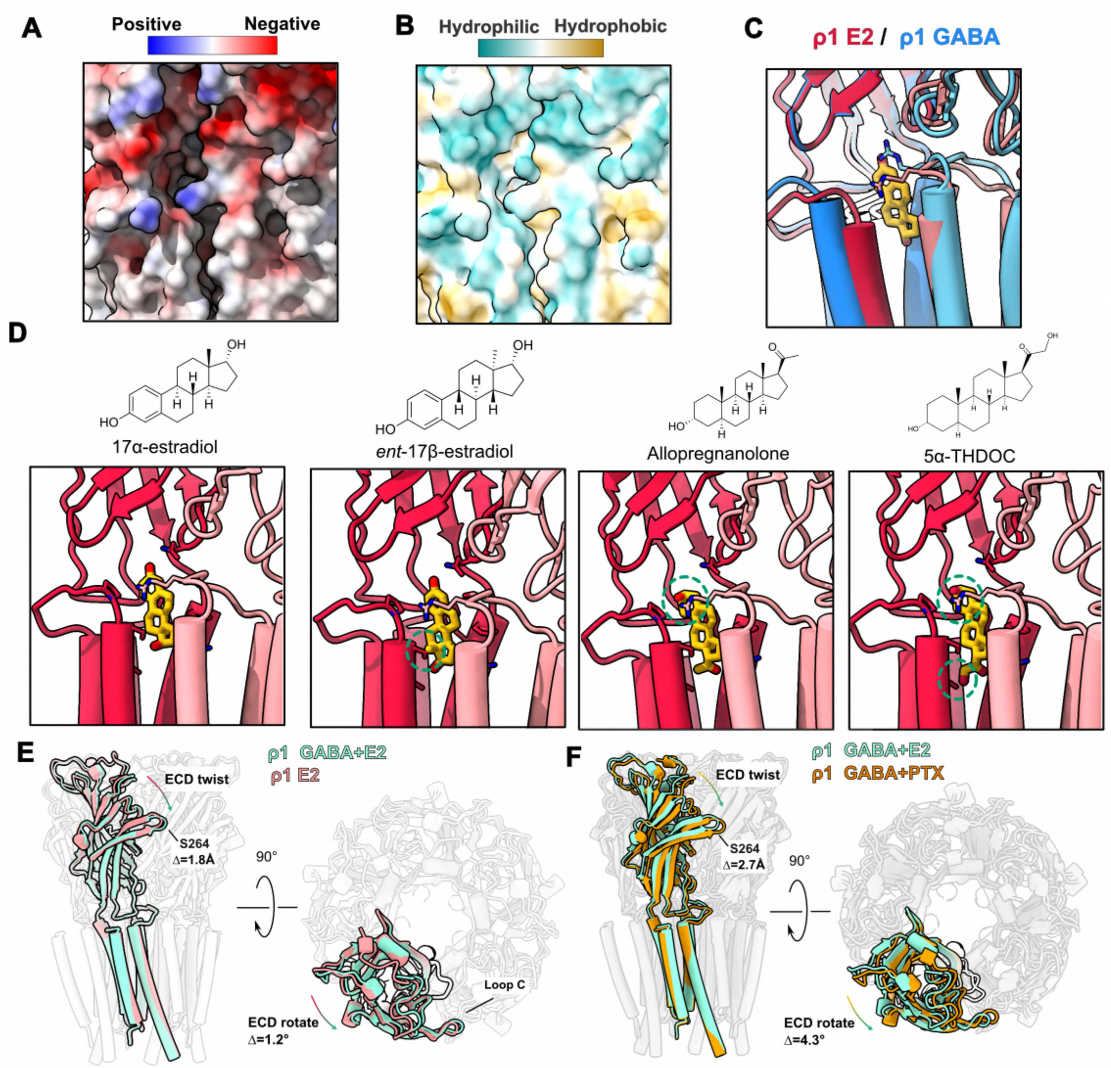


**Figure S4. The binding site of E2.**

(A) Surface representation of a single E2 binding pocket, viewed as in Figure 1F, colored by electrostatic potential.

(B) Surface representation as in A, colored by hydrophobicity.

(C) A single subunit interface as in A–B, showing a superimposition of ρ1-EM structures determined in the presence of E2 (red) or GABA (blue). Conformational changes upon activation/desensitization alter the E2 site, likely precluding binding.

(D) Modeling of E2 homologs 17α-estradiol, ent-17β-estradiol, allopregnanolone and 5α-THDOC into a single interface of the structure with E2. The molecules are manually aligned on the steroid rings of E2. Clashes between the protein and steroids are indicated by the green circles.

(E) Superimposed structures of ρ1-EM with E2 alone (pink) and with E2 and GABA in the primed state (green), viewed from the membrane plane (left) or extracellular side (right). Only one subunit is highlighted for clarity.

(F) Superimposed structures of GABA-bound structures of ρ1-EM with E2 in the primed state (green) and with PTX in the uncoupled intermediate state (gold, PDB ID: 8OQA). Only one subunit is highlighted for clarity.

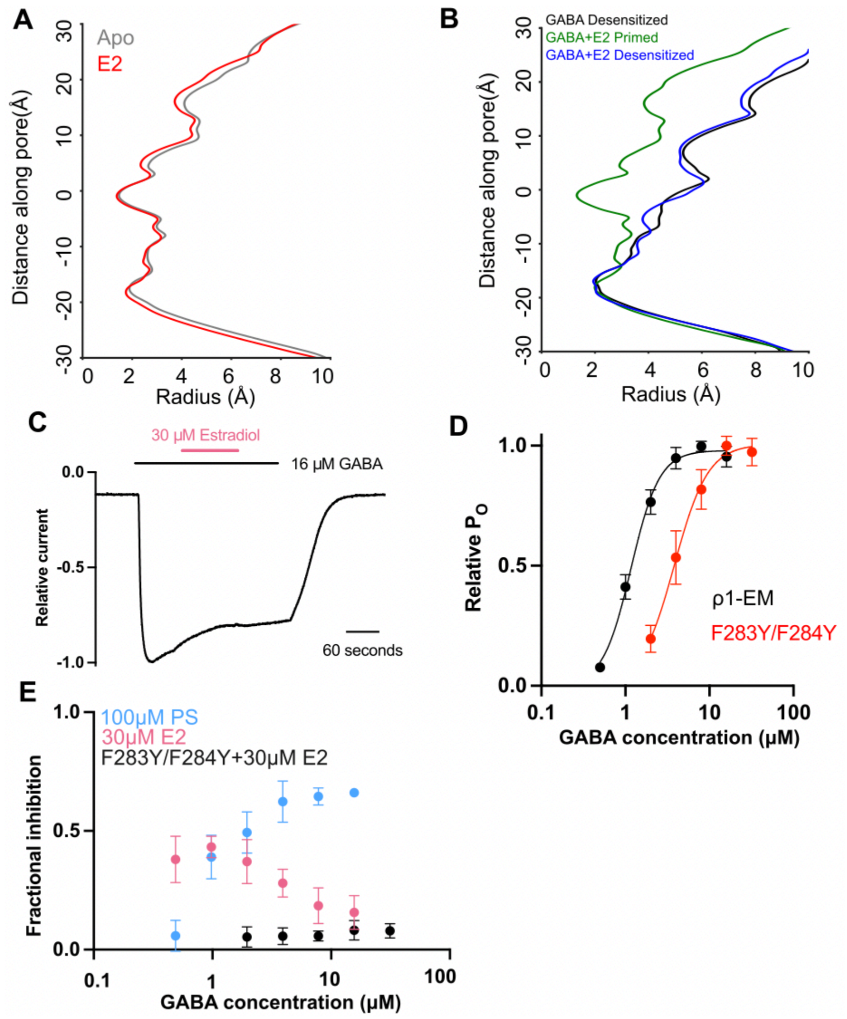


**Figure S5. Pore-radius profiles and electrophysiology of ρ1-EM with E2.**

(A) Pore-radius profiles of ρ1-EM apo (gray) and E2 (red) structures. Both structures are assigned to resting-like states.

(B) Pore-radius profiles of ρ1-EM in the presence of GABA alone (black, PDB ID: 8OP9) and GABA+E2 in the primed (green) and desensitized (blue) states.

(C) Sample trace from TEVC recording of ρ1-EM in response to a saturating GABA concentration (16 μM) alone or in combination with 30 μM E2 as indicated in bars at top of plot.

(D) GABA concentration response curves for ρ1-EM (black) and the F283Y/F284Y mutant (red). Error bars represent SEM for 8 independent oocytes for ρ1-EM or 3 oocytes for the mutant. Solid lines represent fits to Boltzmann curves with an $EC_{50}$ of 1.2 μM (ρ1-EM) or 3.8 μM (F283Y/F244Y).

(E) Fractional inhibition by 30 μM E2 (pink) or 100 μM PS (blue) of the ρ1-EM current response, plotted against co-applied GABA concentrations. Black points represent fractional inhibition of the F283Y/F284Y mutant by 30 μM E2. Error bars represent SEM, derived from 5 independent oocytes for ρ1-EM with 30 μM E2, or 3 independent oocytes for other two conditions.

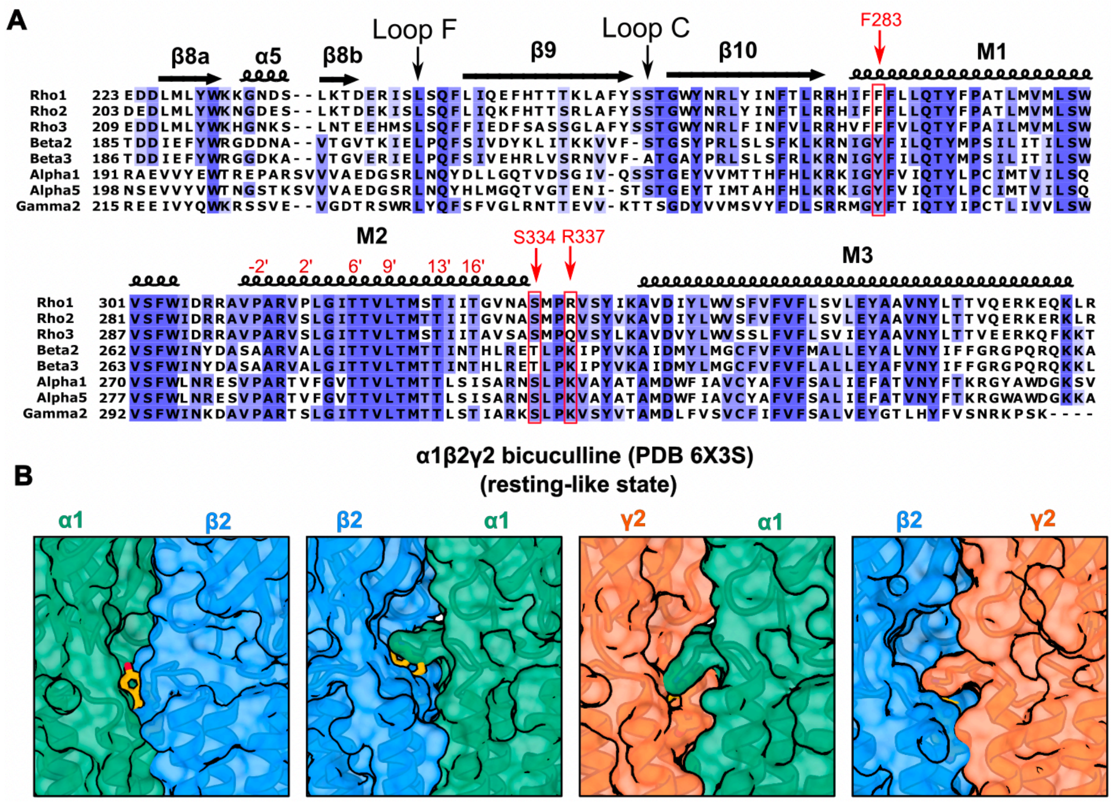


**Figure S6. Sequence alignment of the E2 binding site.**

(A) Sequence alignment of representative human ρ, β, α and γ2 $GABA_A$ receptor subunits. Residues are numbered according to reference UniProt sequences. Key structural features are labeled above alignment, including secondary structure elements and agonist-binding loops C and F. Red labels indicate TMD residues interacting with E2, and M2 pore-facing residues in prime notation. For simplicity, only residues between β8 and M3 relevant to steroid binding in this work are shown.

(B) Surface representations, depicted as in Supplementary Figure S4A, showing zoom views of the four types of subunit interfaces in a previously reported cryo-EM structure of the α1β2γ2 $GABA_A$ receptor (PDB ID: 6X3S). The structure contains the competitive inhibitor bicuculline and is assigned, like ρ1-EM with E2, to a resting-like state. Superimposition of E2 (yellow) from the corresponding ρ1-EM structure indicates little capacity for binding in this site.

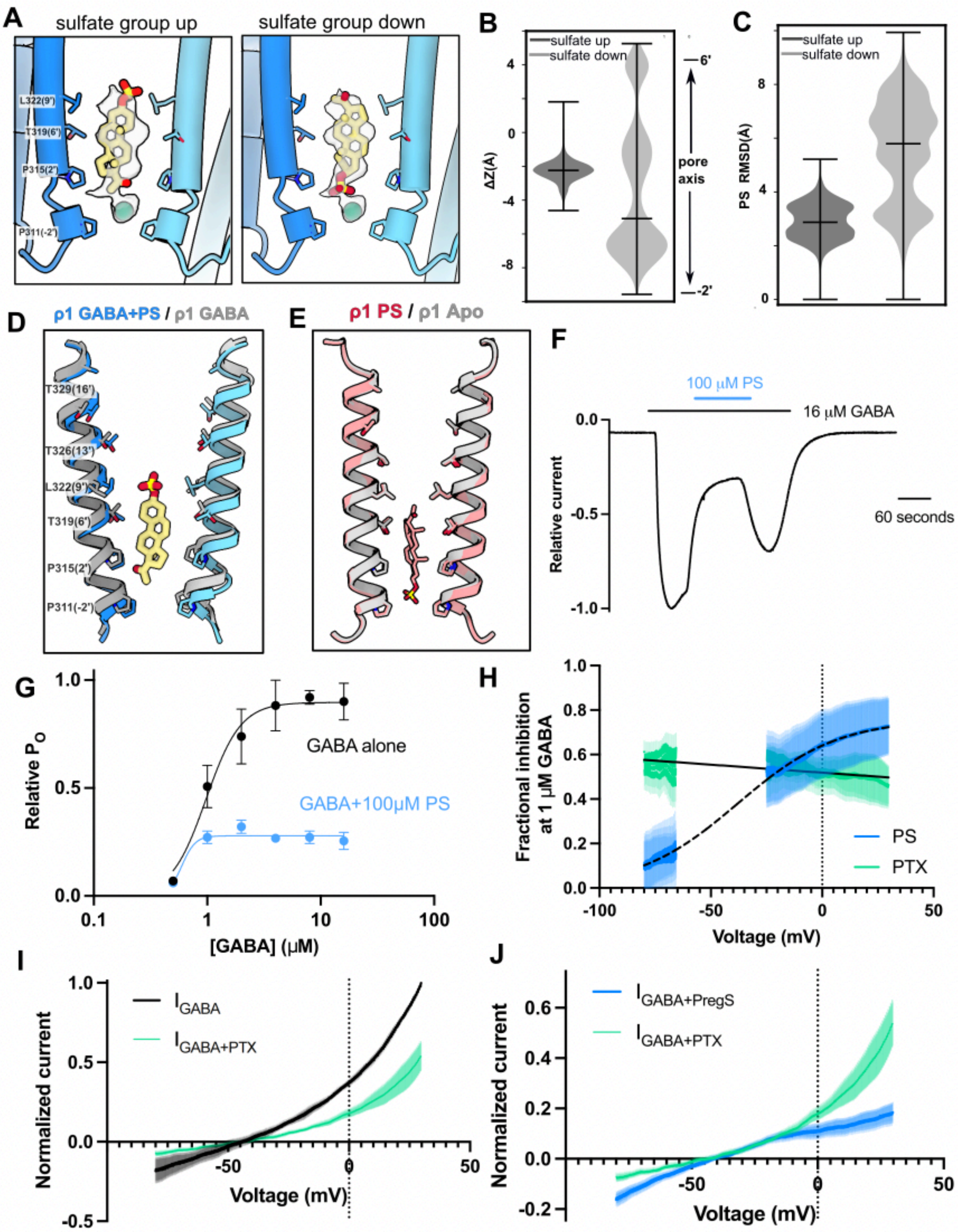
A
sulfate group up
sulfate group down
L322(9')
T319(6')
P315(2')
P311(-2')
B
sulfate up
sulfate down
ΔZ(Å)
pore axis
6'
-2'
C
PS RMSD(Å)
D
ρ1 GABA+PS / ρ1 GABA
T329(16')
T326(13')
E
ρ1 PS / ρ1 Apo
F
100 μM PS
16 μM GABA
Relative current
60 seconds
G
Relative PO
GABA alone
GABA+100μM PS
[GABA] (μM)
H
Fractional inhibition at 1 μM GABA
PS
PTX
Voltage (mV)
I
Normalized current
IGABA
IGABA+PTX
Voltage (mV)
J
Normalized current
IGABA+PregS
IGABA+PTX
Voltage (mV)

**Figure S7. MD simulations and electrophysiology of ρ1-EM with PS.**

(A) Zoom views of the inner pore of ρ1-EM determined with GABA and PS, with experimental density assigned to PS and chloride shown in transparency. Two possible poses are shown for PS, either with the sulfate group oriented up towards the 9' hydrophobic gate (left) or down towards the cytosol (right). PS (yellow), chloride (green) and surrounding residues are shown as sticks and labeled.

(B) Translocation of PS along the pore z-axis in MD simulations launched from the two poses shown in A. Simulation frames are aligned on Cα atoms of the M2 pore-lining helices, and translocation (ΔZ) calculated for the center of mass of PS non-hydrogen atoms along a linear axis passing through the channel pore. Violin plots represent probability densities from 4 independent simulation replicates of >400 ns each, sampled every 0.4 ns (n > 4000), with markers indicating median and extrema.

(C) Mobility of PS in MD simulations as in B, calculated from RMSD of PS non-hydrogen atoms.

(D) Superimposition of two opposing M2 helices in structures of GABA-bound ρ1-EM in the absence (gray, PDB ID: 8OP9) and presence (blue) of PS.

(E) Superimposition of two opposing M2 helices in structures of ρ1-EM in the absence (gray, PDB ID: 8OQ6) and presence (red) of PS.

(F) Sample trace from TEVC recording of ρ1-EM in response to a saturating GABA concentration (16 μM), alone or in combination with 100 μM PS as indicated in bars at top of plot.

(G) GABA concentration response curves in the absence (black) and presence of 100 μM PS (blue). Error bars represent SEM from 5 independent oocytes. Solid lines represent fits to Boltzmann curves with an $EC_{50}$ of 1.0 μM (GABA alone) or 0.6 μM (GABA+PS).

(H) Fractional inhibition of 1 μM GABA responses upon co-application with 100 μM PS (blue) or 500 nM PTX (green), plotted against membrane potential. Shaded region represents SEM for 4 independent oocytes. Points within 15 mV of the reversal potential were removed due to large errors associated with division by values near 0.

(I) Background-subtracted and normalized current-voltage curves for voltage ramps in the presence of GABA alone (black) or in combination with 500 nM PTX (green). Shaded regions represent SEM from 4 independent oocytes.

(J) Overlay of current-voltage responses to GABA co-applied with PTX (green) or PS (blue) normalized to the maximum response to GABA alone.
